# Evolution of ion channels in the water-to-land transition of vertebrates

**DOI:** 10.64898/2026.04.08.717291

**Authors:** Cristóbal Uribe, Gonzalo Riadi, Juan C. Opazo

## Abstract

The transition of vertebrates from aquatic to terrestrial environments represents one of the most profound evolutionary events in their history, involving extensive physiological and morphological innovations. Key adaptations included the transformation of fins into limbs with digits to enable efficient terrestrial locomotion, the ability to perceive novel environmental stimuli, and the emergence of reproductive strategies suited to life on land, processes in which ion channels played fundamental roles. Accordingly, understanding the genetic basis of vertebrate terrestrialization requires investigating the evolution of this group of membrane proteins. Our analyses reveal that the proportion of ion channel genes is highly conserved, representing approximately 1.4% to 1.6% of total protein-coding genes in most lineages, with a notable increase to ∼1.9% in teleost fishes. Our natural selection analyses revealed an overrepresentation of specific ion channel gene families, including TRP, RyR, HTR3, and HCN. We identified 29 ion channel genes showing signatures of positive selection, many of which are associated with key physiological functions such as nociception and thermosensation. We also detected an elevated rate of gene turnover in the common ancestor of terrestrial vertebrates, indicative of substantial genomic remodeling through gene gain and loss. Together, these findings suggest that, despite overall conservation in the proportions of ion channel genes, specific gene families underwent changes that were likely critical to meeting the physiological demands of terrestrial life. These results provide a foundation for future comparative and functional studies aimed at elucidating the molecular mechanisms underlying major environmental transitions.

**Significance:** The transition from water to land was one of the most important events in vertebrate evolution, requiring animals to detect new environmental cues, sense temperature, coordinate movement, and cope with unfamiliar pathogens and physical stresses. Because ion channels control many of these physiological processes, they provide an ideal system for investigating the molecular basis of vertebrate terrestrialization. By tracing their evolutionary history across vertebrates, we show that many ion channels underlying these functions underwent adaptive evolution during the transition from water to land. Our findings suggest that adaptive changes in ion channels helped vertebrates acquire the physiological capabilities needed to survive and diversify on land, providing new insight into the molecular basis of one of the greatest phenotypic transformations in vertebrate evolution.

## Introduction

One of the most fascinating challenges in evolutionary biology is understanding the genetic underpinnings behind phenotypic transformations. Vertebrates have successfully colonized an exceptional range of environments, a diversification that required profound morphological and physiological changes (Ashley-Ross et al., 2013). The growing availability of high-quality genomes now offers an outstanding opportunity to uncover the molecular changes underlying those phenotypic transformations. Well-studied examples include the transition from land to water in cetaceans (whales, dolphins and porpoises) (Nery et al., 2013; Tsagkogeorga et al., 2015; McGowen et al., 2012; Sun et al., 2013), the conquest of the deserts by rodents (Rocha et al., 2021; Yuan et al., 2025), adaptation to subterranean habitats in fossorial mammals (Bondareva et al., 2023; Du et al., 2015; Jiang et al., 2020), and the terrestrialization of invertebrates groups (Martínez-Redondo et al., 2023; Aristide & Fernández, 2023). Among vertebrates, the colonization of land, dated to ∼408-352 million years ago (Kumar et al., 2022), represented one of the most significant events during their evolutionary history (Ashley-Ross et al., 2013; MacIver & Finlay, 2022). This transition was enabled by a suite of morphological and physiological innovations that allowed early vertebrates to meet the challenges of terrestrial life. For example, fins transformed into digit-bearing limbs capable of supporting body weight and locomotion on land, lungs evolved to replace or complement gills for aerial respiration, sensory systems were rewired to detect airborne sounds, odors, and visual cues, and reproductive strategies became less-water dependent, most notably with the origin of the amniote egg (Cupello et al., 2022; Clack, 2009; Clack, 2006).

Ion channels are multimeric proteins embedded in the plasma membrane (Catacuzzeno et al., 2025) that form selective pores, regulating the passage of specific ions (e.g., sodium, potassium, and calcium) across the cell membrane and generating electrical signals that are crucial for nerve signaling and muscle contraction (Barker et al., 2017). These proteins play a key role in the major adaptive challenges associated with colonizing new environments. For example, in locomotion, ion channels such as voltage-gated sodium channels are crucial for generating and propagating action potentials along neurons that initiate movement (Marban et al., 1998). The expansion of voltage-gated sodium channels in early tetrapods is linked to the evolution of new sensory receptors and more complex motor control required for terrestrial locomotion (Zakon et al., 2011; Zakon, 2012). In the sensory system, ion channels also play key roles, TRP channels, for instance, participate in cold, heat, and chemical detection (M. Zhang et al., 2023; Willis, 2009; Saito & Tominaga, 2017; Lu et al., 2022), mechanically-gated ion channels like PIEZO responds to physical force, such as pressure, touch, and sound waves (Delmas & Coste, 2013; Hamed et al., 2024), and cyclic nucleotide-gated (CNG) channels participate in visual and olfactory systems, converting sensory stimuli into electrical signals (Pifferi et al., 2006). Finally, in reproduction, ion channels such as CATSPER (1-4), KCNU1, and members of the TRP channel family participate in fundamental processes of fertility, including sperm capacitation, motility, and the acrosome reaction (Shukla et al., 2012; Dubey et al., 2024; H. Wang et al., 2021). Thus, given their central roles in the emergence of key physiological and morphological traits, reconstructing their evolutionary history is essential for understanding vertebrate evolution.

To study the evolution of ion channels during the colonization of land by vertebrates, we designed a bioinformatic pipeline to annotate the entire repertoire of ion channel genes across the genomes of species representing all major groups of jawed vertebrates. After identifying the ion channel genes in each species, we inferred homologous relationships to investigate the role of positive selection and gene copy number variation. Our main results show that, on average, vertebrates possess a relatively consistent proportion of ion channel genes, ranging from 1.4% to 1.9% of their total protein-coding genes. Furthermore, we identified 29 ion channel genes showing signatures of positive selection that are associated with physiological functions such as nociception, thermosensation, and others. Importantly, we also detected a high rate of gene turnover in the ancestor of terrestrial vertebrates, potentially reflecting adaptive genomic changes associated with life on land. These results highlight that, despite the overall conservation of ion channel gene proportions, specific gene families experienced adaptive changes that were likely crucial for meeting the physiological demands of terrestrial life.

## Results and Discussion

### Annotation and comparative distribution of ion channels in jawed vertebrates

After completing the annotation stage, we found that the 23 sampled mammals possess, on average, 20,868 protein-coding genes, of which approximately 300 (1.4%) encode ion channels subunits. This value is higher than the one we previously estimated for the group (Uribe et al., 2024), a difference likely attributable to changes in the E-value threshold and updates to CDD libraries related to ion channels in version 3.21 (J. Wang et al., 2023). Among sauropsids (birds and reptiles, n=32), the mean number of protein-coding genes was 18,084, with about 287 (1.6%) identified as ion channels subunits (Fig. 3). The eight amphibian genomes contained an average of 21,718 protein-coding genes, with 319 (1.5%) corresponding to ion channels subunits (Fig. 3). In lungfish, we detected 22,039 protein-coding genes, with 329 (1.5%) corresponding to ion channels subunits (Fig. 3), while the coelacanth genome contained 19,737 protein-coding genes, including 332 (1.7%) ion channels subunits (Fig. 3). For the 10 teleost species included in our sampling, the mean number of protein-coding genes was 22,828 genes, with 442 (1.9%) classified as ion channels subunits (Fig. 3). This proportion of ion channel genes is markedly greater than that observed in other jawed vertebrates and is likely attributable to the additional round of whole genome duplication that occurred in the ancestor of teleosts (Amores et al., 1998; Kuraku & Meyer, 2009; Jaillon et al., 2004). Finally, in cartilaginous fish (n=11), we identified an average of 18,812 protein-coding genes, of which 300 (1.6%) correspond to ion channels subunits (Fig. 3). These counts include both pore-forming channel proteins and integral subunits of multimeric channel complexes.

**Figure 1.**
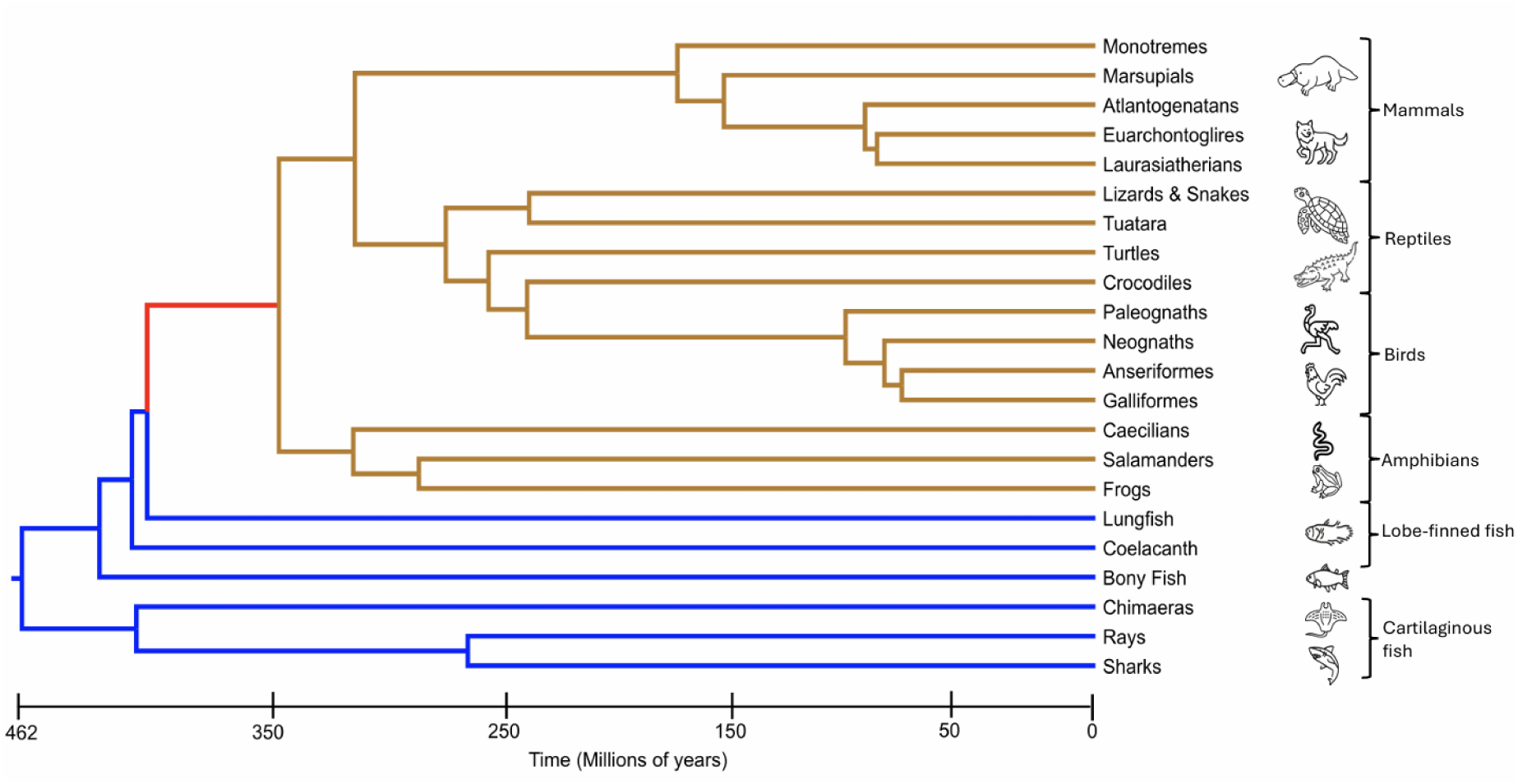
Time-calibrated phylogenetic tree illustrating the jawed vertebrate lineages included in our study. Blue branches represent aquatic lineages, brown branches indicate terrestrial lineages, and the red branch highlights the ancestral lineage that underwent the transition from aquatic to terrestrial environments. Animal silhouettes were obtained from Flaticon (www.flaticon.com), and evolutionary relationships and divergence times were derived from the timetree database (https://timetree.org/).

**Figure 2.**
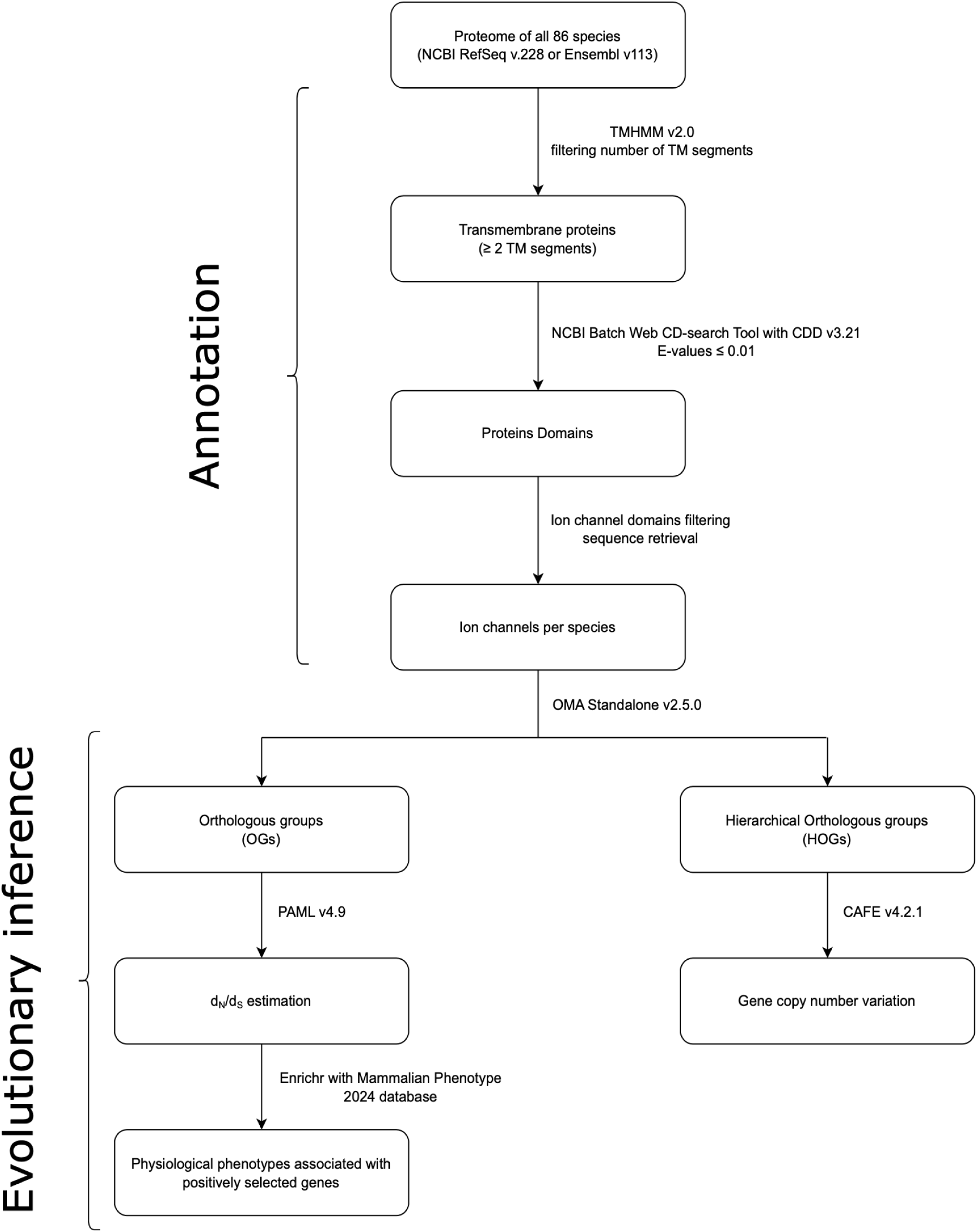
Workflow summarizing the annotation and evolutionary inference pipeline used in this study. TM: Transmembrane; d_N_/d_S_: ratio of non-synonymous to synonymous substitution rates.

**Figure 3.**
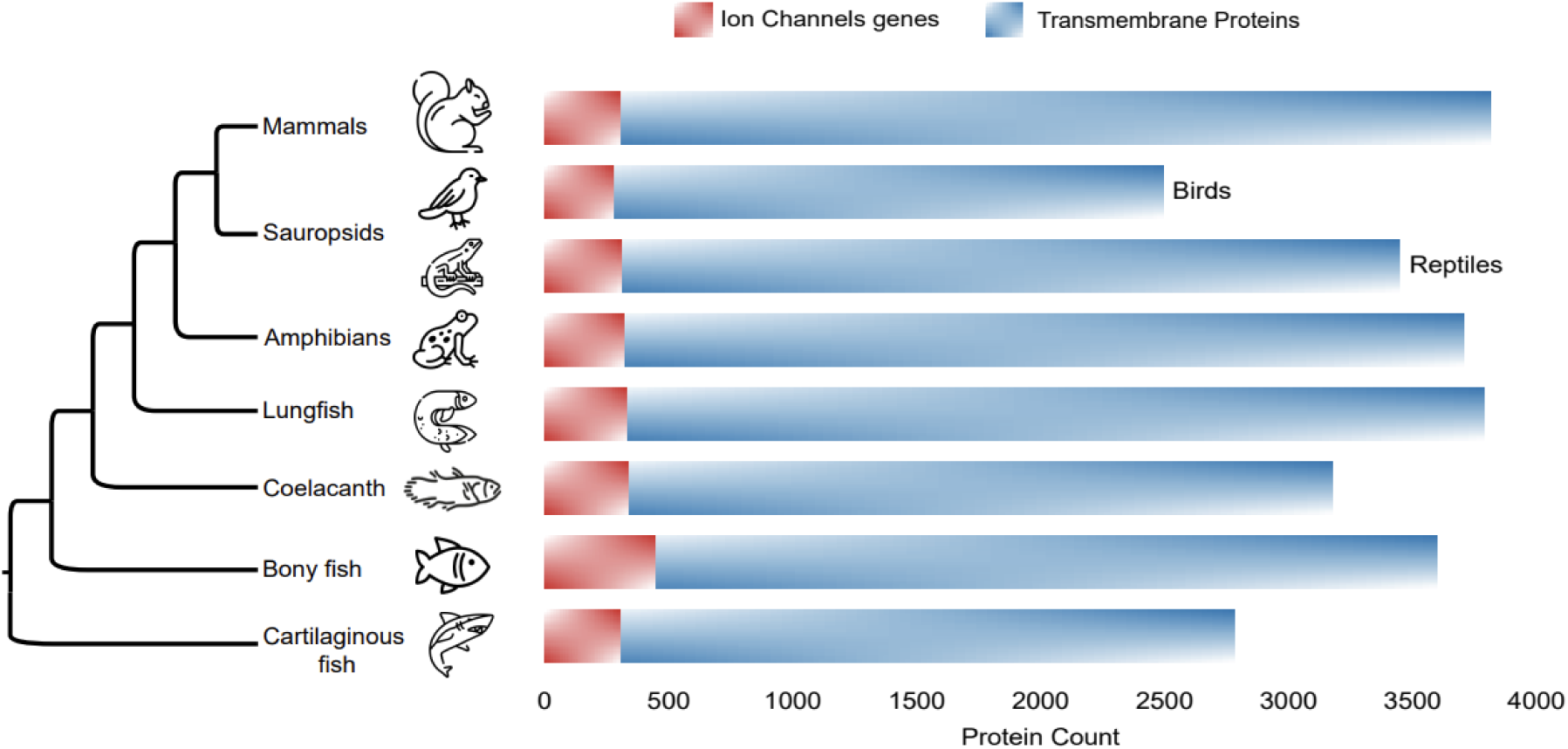
Ion channel and transmembrane protein abundance across jawed vertebrates.

Consistent with previous reports (Hughes & Friedman, 2008; Lovell et al., 2014; G. Zhang et al., 2014), we observed that birds possess fewer protein-coding genes than other vertebrate groups, with values ranging from 14,765 in the common swift (*Apus apus*) to 18,018 in the chicken (*Gallus gallus*). This reduction also extends to transmembrane proteins, which range from 2,032 in the white-throated tinamou (*Tinamus guttatus*) to 3,358 in the North Island brown kiwi (*Apteryx mantelli mantelli*). These findings are consistent with the reduced genome size of birds, which is hypothesized to increase the cell surface relative to volume, thereby maximizing gas exchange and facilitating the metabolic demands of powered flight, a pattern also observed in bats (L. Wu et al., 2021; Gregory, 2002; Gregory, 2001). Despite this overall reduction, the relative abundance of ion channel genes remains stable at approximately 1.7%, comparable to other jawed vertebrates (1.4% to 1.9%), highlighting the essential role of these proteins in maintaining life.

Thus, overall, our comparative analyses show that the relative abundance of ion channel genes varies little among jawed vertebrate lineages, highlighting their fundamental and evolutionarily conserved role in vertebrate physiology.

### Overrepresentation of RyR, HCN, HTR3, and TRP gene families under positive selection

We analyzed a final set of 167 1:1 orthologous groups, each comprising between 11 to 86 species per group (Fig. 4). Notably, over 95% of these groups included more than 30 species (Fig. 4). Within this dataset, we identified 29 ion channel genes exhibiting signatures of positive selection (Supplementary Table S1), suggesting potential adaptive changes associated with the water-to-land transition. Importantly, these genes do not necessarily correspond to independent ion channels, as they encode proteins that either form the ion-conducting pore or constitute integral subunits of multimeric channel complexes.

**Figure 4.**
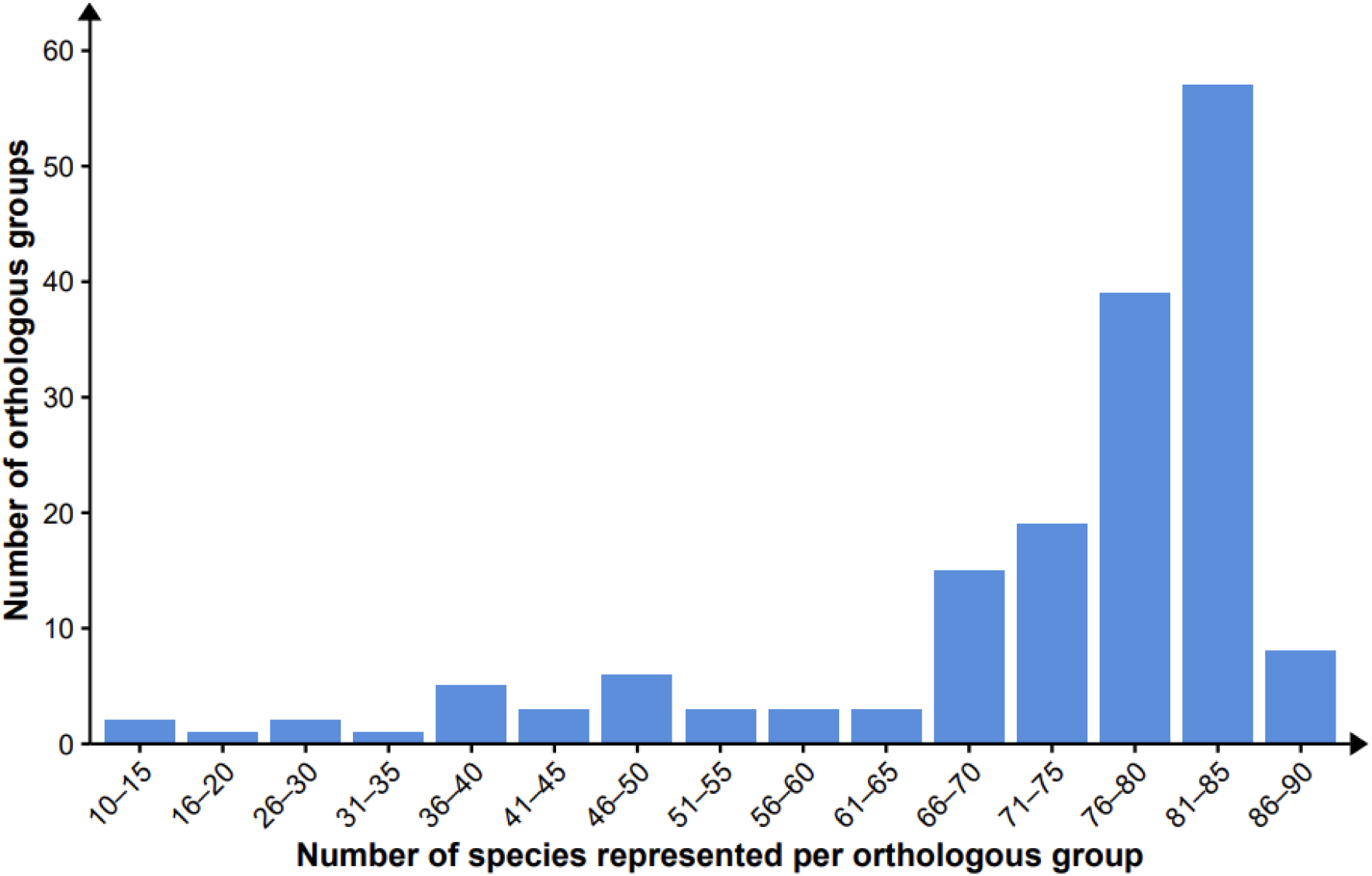
Distribution of orthologous groups according to the number of species they contain.

Our next step was to assess whether the 29 ion channel genes showing the signature of natural selection were non-randomly distributed among gene families. To this end, we first classified each of the 167 analyzed ion channels genes into their respective gene families (Supplementary Table S2) and then compared the observed distribution of the 29 positively selected ion channels genes against their expected family level counts (Fig. 5). Our analysis revealed that ryanodine receptors (RyR), hyperpolarization-activated cation channels (HCN), 5-hydroxytryptamine receptor 3 (HTR3) and transient receptor potential channels (TRP) families accumulated more positively selected genes than expected, with observed/expected ratios ranging from 2.3 to 3.8 (Fig. 5). In contrast, families such as TMEM16, CHRN, CLC, and KCN were under-represented with ratios below 1 (Fig. 5).

**Figure 5.**
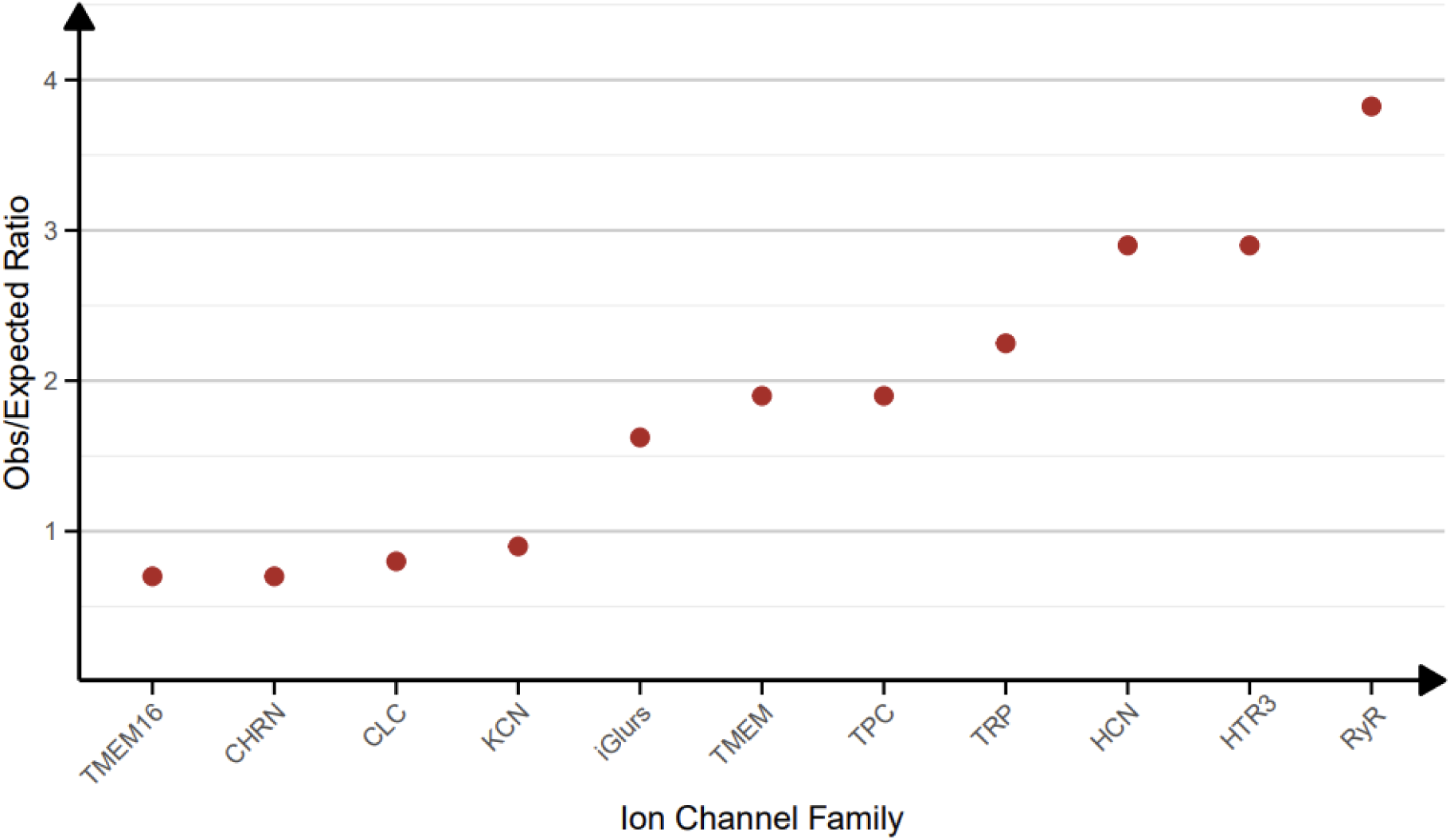
Enrichment analysis of ion channel gene families under positive selection. The plot shows the observed-to-expected ratio for each ion channel family among the 29 genes exhibiting signatures of positive selection.

The most represented ion channel families perform key physiological functions in vertebrates. For instance, RyRs regulate intracellular calcium levels and are essential for excitation-contraction coupling in both cardiac and skeletal muscle cells, as well as for diverse processes including development, signaling, learning, and memory (Lanner et al., 2010). By increasing cytoplasmic calcium concentration, RyRs trigger muscle contractions that sustain coordinated movement and normal cardiac rhythm (H.-T. Yang et al., 2002; Chagovetz et al., 2019). HCN channels are primarily expressed in the heart and the central nervous system, where they play fundamental roles in regulating cardiac and neuronal rhythmicity by generating the pacemaker current (Wahl-Schott & Biel, 2009). This inward cation current is activated during membrane hyperpolarization and carries both sodium and potassium ions, contributing to the depolarization process that underlies the rhythmic beating of the heart (Scicchitano et al., 2012). In the central nervous system, HCN channels are involved in critical functions including learning, memory, and motor control (Z. Zhang et al., 2024). HTR3 channels are expressed in both the central and peripheral nervous systems, where they mediate fast synaptic signaling by opening in response to the neurotransmitter serotonin (Barnes et al., 2009). By allowing the influx of cations such as Na⁺ and Ca^2^⁺, HTR3 channels depolarize postsynaptic neurons and participate in diverse physiological processes, including pain sensation, motor regulation, anxiety control, and emesis (Thompson & Lummis, 2006). The last highly represented group, TRP ion channels, constitutes a large and diverse family of non-selective cation channels expressed in many tissues, particularly in sensory neurons (Nelson et al., 2011; Kauer & Gibson, 2009). They serve as key molecular sensors, detecting a broad spectrum of environmental and physiological stimuli, including temperature, pain, mechanical stress, phototransduction, and chemical irritants (M. Zhang et al., 2023). Owing to their central roles in these essential processes, TRP channels have been widely studied across fields ranging from evolutionary biology to medicine (Saito et al., 2011; Ribelayga, 2010; Yue et al., 2015; Lee et al., 2021; L. Yue & Xu, 2021; Cabezas-Bratesco et al., 2022; Flores-Aldama et al., 2020; Cancino et al., 2025). Together, these findings suggest that adaptive molecular evolution during the water-to-land transition was mainly concentrated in ion channel gene families central to excitability, sensory perception, and the coordination of vital physiological processes.

### Functional enrichment of positively selected ion channels during the vertebrate water-to-land transition

We next used the 29 ion channel genes showing signatures of positive selection as input to an enrichment analysis using the Enrichr tool to gain a more detailed understanding of the biological functions potentially affected during the vertebrate water-to-land transition (Supplementary Table S3). The resulting enriched phenotypic associations clustered into four major physiological categories: 1) neurological system, 2) sensory perception, 3) immune system, and 4) lethality (Fig. 6). Each physiological category is composed of a set of physiological features described by functional or characteristic terms. For example, premature death is a physiological feature of the lethality category.

**Figure 6.**
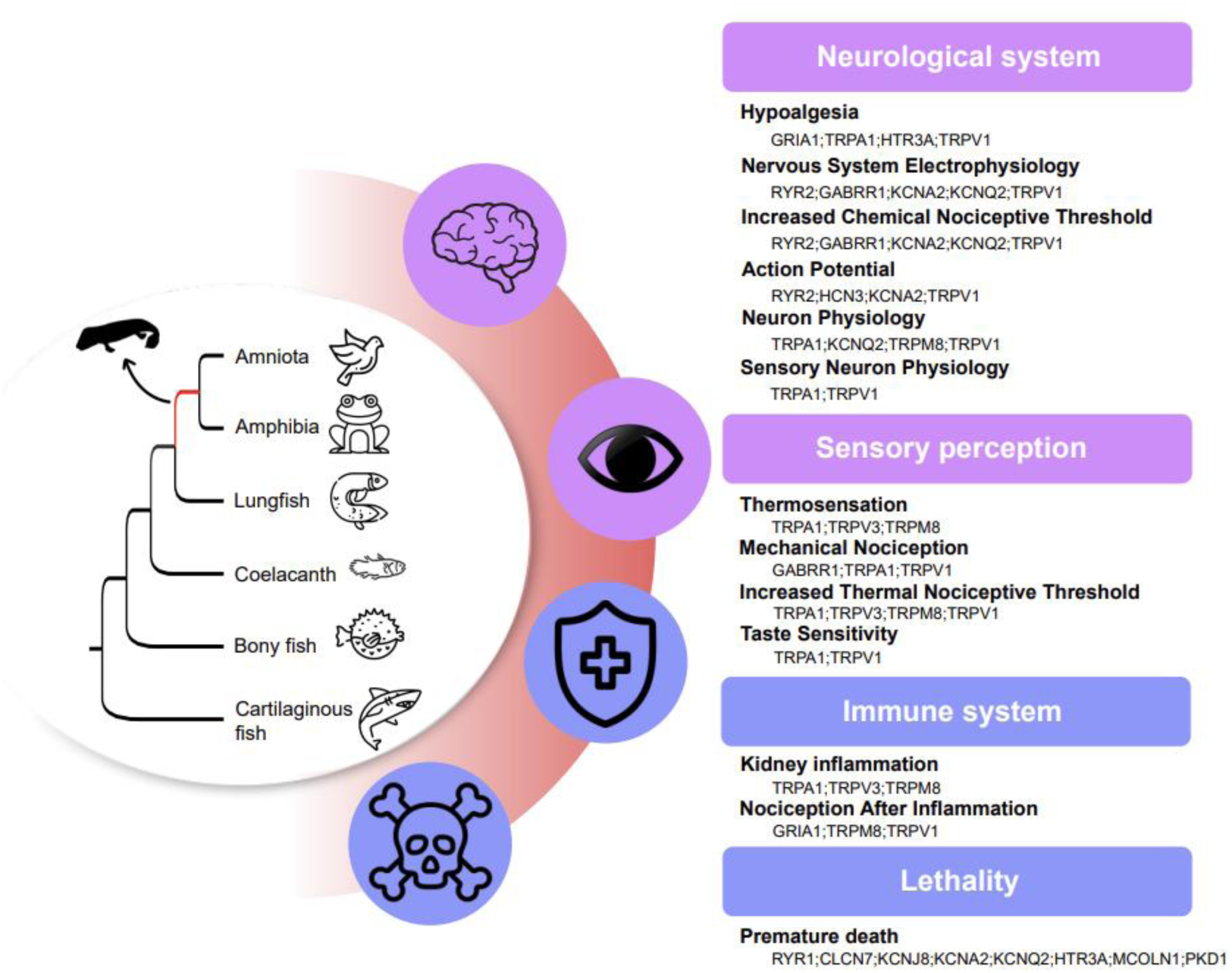
Enrichment of functional categories among ion channel genes under positive selection in the tetrapod ancestor.

Several ion channels were recurrently associated with multiple physiological features across these categories (Fig. 6), suggesting that a subset of ion channels may have played central roles in key adaptive processes during vertebrate evolution.

The physiological features associated with the neurological system category point to evolutionary changes in neuronal signaling and excitability, suggesting that adaptations in the nervous system were a key component of vertebrate terrestrialization. Ion channels most frequently linked to this category included RYR2, GABRR1, KCNA2, and KCNQ2. RYR2 encodes a calcium release channel located in the sarcoplasmic reticulum, essential for excitation-contraction coupling in cardiac muscle (Kobayashi et al., 2022). A comparative study shows that RYR2 exhibits distinct thermal sensitivities between endothermic and ectothermic vertebrates (Vornanen, 2006). In cold water fish, RYR2 maintains robust gating and calcium sensitivity even at temperatures as low as 2°C, whereas in mammals, gating slows and closure is impaired under similar conditions, indicating reduced function under cold stress. These differences highlight lineage-specific adaptations of RYR2 to thermal environments. GABRR1 encodes the rho-1 subunit of the ionotropic GABA receptor, a chloride channel expressed predominantly in the central nervous system and retina, where it contributes to inhibitory neurotransmission (Zhu et al., 2019). In the retina, these channels modulate glutamate release at bipolar cell terminals, thereby shaping the duration of visual signals (Popova, 2014). Such fine-tuning of visual inhibition may have aided adaptation to variable light environments during vertebrate evolution. KCNA2, a member of the voltage-gated potassium channel family KCNA, plays a pivotal role in neuronal excitability and neurotransmitter release (Hoegg & Meyer, 2007). In the cerebellum, Kv1.2 subunits encoded by KCNA2 regulate calcium spiking in Purkinje-cell dendrites and control GABA release from basket-cell terminals. Mutations in KCNA2 in mice cause chronic motor incoordination due to disrupted excitation-inhibition balance (Xie et al., 2010). These functions are essential for motor coordination, a capability critical for efficient terrestrial locomotion. Similarly, KCNQ2 encodes a potassium channel primarily expressed in the nervous system, where it regulates neuronal excitability and helps maintain membrane potential stability via M-current (Devaux et al., 2004; Ma et al., 2023). A mutation in KCNQ2 produces developmental and epileptic encephalopathy, characterized by severe early-onset seizures (Ye et al., 2023; Brun et al., 2022; M. Mao et al., 2025), underscoring its critical role in maintaining neuronal homeostasis.

The sensory perception category was strongly represented by members of the TRP family, particularly the thermoTRP channels TRPV1, TRPA1, TRPM8, and TRPV3, which were associated with multiple physiological features (Fig. 6). These TRP ion channels are multimodal receptors, capable of being activated by temperature changes and a variety of sensory stimuli (Saito & Tominaga, 2017). TRPV1 is activated by high temperatures (>43°C), low pH, and capsaicin (Luo et al., 2019). In contrast, TRPM8 responds to low temperatures (<25°C) and compounds such as menthol and eucalyptol (Palchevskyi et al., 2023). TRPV3 is activated by warm temperatures (33-39°C) and natural compounds such as genariol and carvacrol (Kalinovskii et al., 2023). Lastly, TRPA1 is activated by noxious cold and chemical irritants, including cinnamaldehyde, menthol, and other electrophilic compounds (Kim et al., 2024). Those thermoTRP channels have been extensively studied for their evolutionary significance in vertebrate sensory systems (Lu et al., 2022; Saito & Tominaga, 2017; Laursen et al., 2016; Patapoutian et al., 2003; Vay et al., 2012; Kashio & Tominaga, 2022; Castillo et al., 2018). For example, comparative studies on TRPV1 reveal that thermosensitivity varies among species according to their environmental niches (Saito & Saito, 2025). In squirrels and camels, which tolerate high temperatures, TRPV1 exhibits a higher activation threshold, thereby reducing nociceptive responses under heat stress (Laursen et al., 2016). Such adjustments likely prevent unnecessary pain signaling in persistently hot habitats while preserving the ability to detect damaging thermal extremes. Similarly, variation in TRPV1 and TRPA1 thermosensitivity among *Xenopus* frogs has been linked to differences in habitat temperature, suggesting a role for these channels in thermal niche specialization (Saito et al., 2019; Saito et al., 2016). In addition, the formation of the MHR1-3 domain in TRPM8 confers cold sensitivity, enabling vertebrates, particularly terrestrial tetrapods, to detect potentially harmful cold and trigger behavioral and physiological responses (Lu et al., 2022). Finally, TRPV3 is abundantly expressed in epidermal keratinocytes (Lei & Tominaga, 2025) and exhibits opposite temperature sensitivities across vertebrate lineages. In mammals, it is activated by warm temperatures, whereas in the western clawed frog, it responds to cold stimuli, illustrating that TRPV3 function diverged during vertebrate evolution (Saito et al., 2011). Collectively, these findings indicate that functional diversification within TRP channels paralleled the ecological shifts of vertebrates, facilitating their adaptation to a wide range of thermal niches during the water-to-land transition.

Additional enriched physiological categories included immune system and lethality, encompassing terms such as kidney inflammation, nociception after inflammation, and premature death (Fig. 6). Notably, TRP channels were prominent among the immune-related phenotypic terms (Fig. 6). These channels play key roles in inflammatory processes, acting in both pro- and anti-inflammatory pathways (Silverman et al., 2020; Gouin et al., 2017; Straub, 2014; C.-K. Wu et al., 2021). For example, TRPV1 and TRPA1, expressed in nociceptive neurons and immune cells, are activated by endogenous activators, such as acidified pH, prostaglandins, and oxidative aldehydes, all of which are produced during inflammation (Csekő et al., 2019). TRPV1 promotes neurogenic inflammation by triggering the release of proinflammatory neuropeptides (substance P and CGRP), whereas TRPA1 detects oxidative stress, contributing to chronic inflammatory pain (Straub, 2014). In contrast, TRPM8 exerts anti-inflammatory effects by counteracting TRPV1/TRPA1-mediated hyperexcitability, reducing neuropeptide release, and attenuating edema (Straub, 2014; Silverman et al., 2020; Premkumar, 2014). TRPV3, mainly expressed in keratinocytes, is activated by warm temperatures and endogenous lipids, and contributes to skin inflammation by regulating the release of interleukin-1α, prostaglandin E_2_, and nitric oxide (S. M. Huang et al., 2008; Mandadi et al., 2009). GRIA1, which encodes a subunit of AMPA receptors, mediates fast excitatory transmission in the vertebrate central nervous system and contributes to synaptic plasticity and inflammatory pain via calcium-permeable AMPA receptor upregulation in the dorsal horn (Y.-C. Chen et al., 2001; Hartmann et al., 2004). Together, these interactions between ion channels modulate the balance between nociceptive sensitization and inflammation resolution, supporting vertebrate adaptation to new terrestrial challenges, including exposure to new pathogens, toxins, and physical stressors.

Finally, the lethality category, defined as death after weaning but before the normal life span (Baldarelli et al., 2024), included multiple ion channels whose mutations disrupt vital physiological processes. For example, RYR1 mutations cause congenital myopathies leading to respiratory failure and premature death (Amburgey et al., 2013; O’Connor et al., 2023). CLCN7 mutations cause osteopetrosis, a severe, life-threatening disorder characterized by bone fragility and impaired hematopoiesis (X. Wang et al., 2023). KCNJ8 mutations are associated with cardiac channelopathies such as J-wave syndrome (JWS), which manifests as abnormal ECG patterns and lethal ventricular arrhythmias that can lead to sudden cardiac death (Medeiros-Domingo et al., 2010). Other mutations have been linked to sudden infant death syndrome (SIDS), particularly under conditions of infection or inflammation, reflecting its critical role in regulating vascular tone and the cardiac adaptive response to stress (Tester et al., 2011). Additionally, mutations in this channel contribute to Cantu syndrome, a multi-organ disease characterized by vascular abnormalities that may increase the risk of infant mortality (Nichols, 2016). Although mutations in KCNA2 are not linked to a defined human disease, they are associated with severe seizure disorders, and knockout experiments in mice have demonstrated premature death (Syrbe et al., 2015). Similarly, mutations in KCNQ2 cause a spectrum of developmental and epileptic encephalopathies (DEEs), including early infantile epileptic encephalopathies (EIEEs), which are marked by high mortality and morbidity during infancy (D. Y. Chen et al., 2018). HTR3A is a ligand-gated ion channel that mediates excitatory responses to serotonin in the nervous system. Mutations in this channel have been shown to cause premature death in mice from complications related to bladder overactivity, chronic urinary retention, and urinary tract outlet obstruction (Bhattacharya et al., 2004). For MCOLN1, a cation channel present in the membranes of lysosomes and endosomes that plays a fundamental role in the ability of cells to digest and recycle materials, mutations can cause mucolipidosis type IV (MLIV). MLIV is a lysosomal storage disease that exhibits progressive intellectual disability, motor impairment, and premature death (Boudewyn et al., 2017). Finally, PKD1 is an integral membrane protein that participates in diverse processes, including regulation of cell growth, division, and motility, as well as interactions with other cells, particularly in the kidney and liver. Mutations in PKD1 in Autosomal Dominant Polycystic Kidney Disease (ADPKD) can cause vascular abnormalities, particularly ruptured intracranial aneurysms, leading to serious health problems such as stroke, brain damage, coma, and even death (Rossetti et al., 2003). Taken together, these examples illustrate that mutations in ion channels, despite their functional and structural diversity, consistently disrupt core physiological processes essential for survival. Their recurrent association with premature death underscores the fundamental role of ion channels in maintaining organismal viability and highlights why they have been key targets of selective pressures during the vertebrate water-to-land transition.

### Accelerated gene turnover rate in the tetrapod ancestor

In our copy number variation (CNV) analysis, we observed variation in gene turnover rates (Fig. 7). In the first model, we estimated two λ parameters, one for terrestrial and another for aquatic vertebrates. This analysis revealed that aquatic vertebrates exhibit a higher gene turnover rate (λ = 0.00099), which is 1.45 times faster than that of terrestrial vertebrates (λ = 0.00068). In the second model, we estimated three λ parameters, one for terrestrial vertebrates, one for aquatic vertebrates, and a third for the tetrapod ancestor (Fig. 7). Consistent with our two-λ model, aquatic vertebrates showed a higher evolutionary rate (λ = 0.00099), 1.47 times faster, than terrestrial vertebrates (λ = 0.00067). However, the highest λ parameter was observed in the last common ancestor of tetrapods (λ = 0.00123), which is 1.83 times higher than that of terrestrial vertebrates (λ = 0.00067) and 1.24 times higher than that of aquatic vertebrates (λ = 0.00099). The faster gene turnover rate in aquatic vertebrates could be influenced by the whole genome duplication (WGD) that occurred in the common ancestor of all extant teleosts around 300 million years ago (Amores et al., 1998; Kuraku & Meyer, 2009; Jaillon et al., 2004).

**Figure 7.**
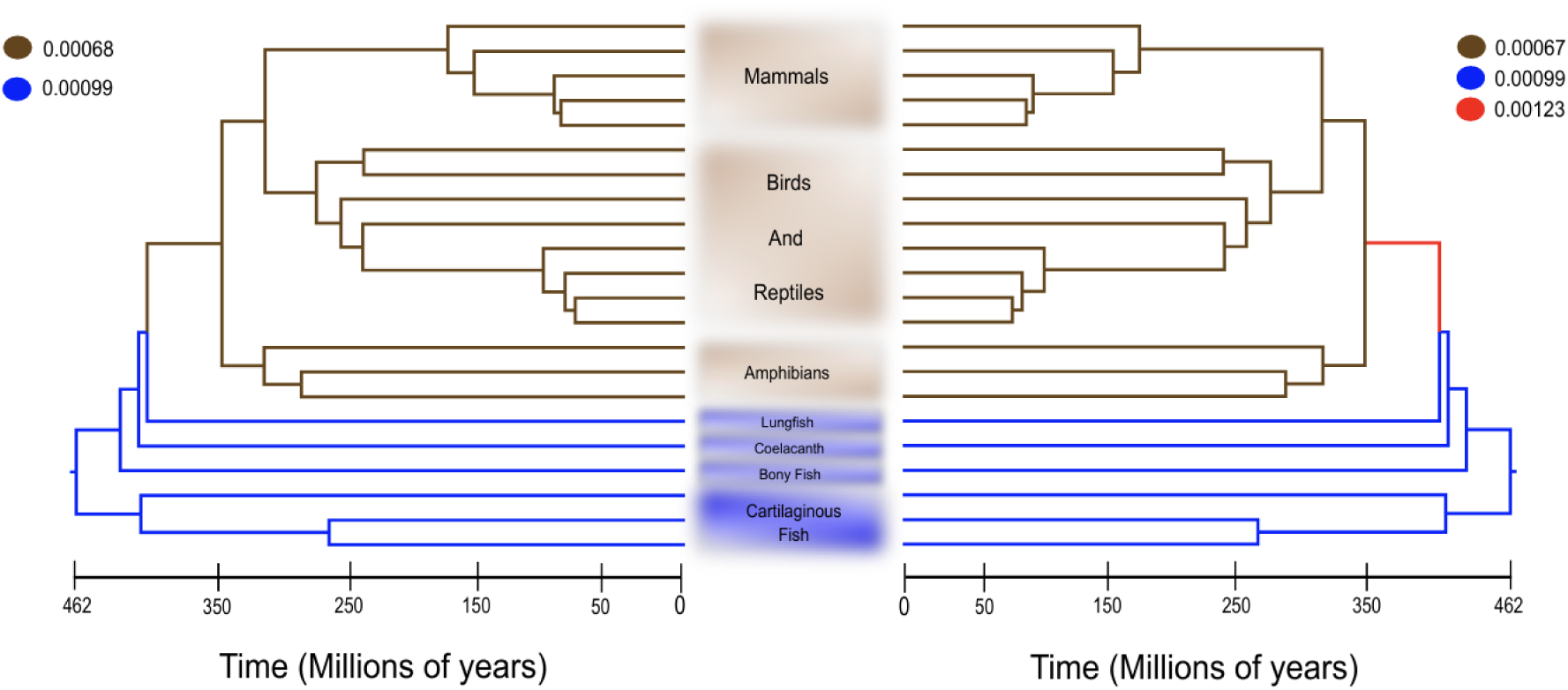
Estimated gene turnover rates of ion channels across jawed vertebrates. The first model (left) estimates separate gene turnover rates (λ) for ion channels in terrestrial vertebrates (brown branches) and aquatic vertebrates (blue branches). The second model (right) estimates separate λ values for terrestrial vertebrates (brown branches), aquatic vertebrates (blue branches), and the last common ancestor of tetrapods (red branch).

CNV analysis examines differences in gene copy number across the genomes of sampled species along the phylogeny. This type of genetic variation represents a significant source of genetic diversity (F. Zhang et al., 2009). CNVs can shape phenotypic traits, influence adaptation to novel environments, and affect susceptibility to diseases, among other biological processes (Bhanuprakash et al., 2018). For example, the expansion of the amylase gene (AMY), which enables starch digestion, has been reported in several mammalian species that consume starch-rich diets, such as humans, mice, rats, pigs, and dogs (Pajic et al., 2019). Moreover, an intraspecific correlation has been observed in humans, with populations that historically consume more starch tending to have higher copy numbers of amylase genes (Perry et al., 2007). Another well-studied example is found in Antarctic icefishes (*Channichthyidae*), in which CNVs have shaped extreme physiological adaptations (Corliss et al., 2019). These species have lost hemoglobin and myoglobin genes, resulting in their “white blood”, while a novel antifreeze glycoprotein gene (AFGP) arose through the duplication and neofunctionalization of a digestive enzyme gene (L. Chen et al., 1997). Together, gene loss and duplication allowed these species to survive in subzero waters, underscoring the role of CNVs in vertebrate adaptation (Shin et al., 2024).

Together, these results indicate that the highest gene turnover occurred in the last common ancestor of tetrapods, suggesting an episode of intensified gene gain and loss during this evolutionary transition. This elevated λ contrast with the lower rates observed in both extant terrestrial and aquatic lineages, highlighting the tetrapod ancestor as a key period of genomic innovation. Such a pattern is consistent with the major ecological and physiological shifts associated with the water-to-land transition, where extensive remodeling of gene repertoires, including ion channels, may have been required.

## Conclusions

In this study, we investigated the evolutionary dynamics of ion channels during the vertebrate transition from water to land, which occurred approximately 408–352 million years ago. It is well known that this major ecological shift required extensive physiological reorganization, involving processes in which ion channels play central roles, including air breathing, osmoregulation, terrestrial locomotion, neuromuscular control, sensory transduction, and host–pathogen interactions. Interestingly, despite surveying a broad diversity of jawed vertebrate lineages, we found that the relative proportion of ion channel genes is remarkably conserved (∼1.4–1.9%), highlighting their fundamental role in vertebrate physiology. Our natural selection analyses revealed an overrepresentation of specific ion channel gene families, including TRP, RyR, HTR3, and HCN. Genes under positive selection were predominantly associated with the nervous and immune systems, sensory perception, and essential viability traits, and were implicated in key physiological processes, including thermosensation, pathogen response, and nociception. Our findings suggest that excitability and sensory pathways underwent substantial remodeling during vertebrate terrestrialization. Importantly, our study focuses on genes encoding ion channel subunits, which constitute the molecular components of functional ion channels. Therefore, our results reflect the evolutionary dynamics of these subunits rather than the properties of fully assembled channel complexes. Finally, our results indicate that gene copy number variation also played a pivotal role in this transition, as evidenced by the highest estimated λ value occurring in the common ancestor of terrestrial vertebrates.

## Material and Methods

### Taxonomic sampling, DNA sequences, and ion channel annotation

We assembled a dataset of 86 species spanning all major groups of jawed vertebrates. Our sampling included cartilaginous fishes, bony fishes, coelacanths, lungfish, amphibians, reptiles, birds, and mammals (Fig. 1). Species were chosen from genome databases using strict quality filters. We included only assemblies deposited in NCBI RefSeq v.228 (Sayers et al., 2025), assembled at the chromosome level, and sequenced at ≥30x coverage. With a few exceptions, all 86 selected species met these criteria (Supplementary File S1).

After selecting the species, we downloaded their protein-coding sequences (.faa file extension) from the Ensembl v.113 (Harrison et al., 2024) and NCBI RefSeq v.228 (Sayers et al., 2025) databases. For each gene, only the longest transcript isoform was retained. We then predicted transmembrane segments with TMHMM v2.0 (Krogh et al., 2001) and kept proteins with ≥2 transmembrane domains. Conserved domains were annotated using the NCBI Batch Web CD-search Tool (Marchler-Bauer et al., 2011) against the CDD database v.3.21 (J. Wang et al., 2023) with an E-value threshold of 0.01. To identify ion channels, we compiled a list of ion channel-associated CDD entries and intersected it with the CD-search results using an in-house Perl script, yielding a preliminary set of ion channels per species. Finally, we manually curated these sets, consulting NCBI and GeneCards (Stelzer et al., 2016) to remove putative false positives and produce the final per-species ion channel lists (Ion_channel_per_species_list.zip), which were used in downstream analysis (Fig. 2).

### Homology inference

Once the ion channels genes were identified for each species, we inferred homologous relationships using OMA Standalone v2.5 (Altenhoff et al., 2019), which identified Orthologous Groups (OGs) and Hierarchical Orthologous Groups (HOGs). To maximize the number of OGs available for analysis, we employed a non-exact search strategy (Nery et al., 2013). In this procedure, we specified the species/clades that must be represented in each orthologous group, as well as the minimum number of species per orthologous group. Specifically, we required that OGs include at least one amniote, the lungfish, one amphibian, and one bony fish, with a minimum of 11 species per orthologous group. This approach ensured an appropriate phylogenetic coverage while maintaining sufficient data density for robust comparative analyses. Using the OGs files as input, we estimated the ratio of non-synonymous (d_N_) to synonymous (d_S_) substitution rates to assess the role of natural selection. The HOGs files, in turn, were used to analyze gene copy-number variation. For the d_N_/d_S_ analysis, amino acid sequences were aligned with MAFFT v7.490 (Katoh & ^S^_tandley, 2013_^)^ with the FFT-NS-2 strategy, and codon-based nucleotide alignments were obtained by employing the amino acid alignments as templates, using the software PAL2NAL (Suyama et al., 2006).

### Molecular evolution analyses

Codon-aligned nucleotide sequences generated with PAL2NAL were used as input for PAML v4.9 software (Z. Yang, 2007) in Phylip format. We estimated the ratio of non-synonymous (d_N_) to synonymous (d_S_) substitution rates (ω = d_N_/d_S_) using the branch-site model approach. This approach compares two nested models. In the first (alternative) model, the branches of interest, called foreground branches, are allowed to have a class of sites evolving under positive selection (ω_2_ > 1), while the remaining branches, called background branches, are modeled with two site classes: conserved sites with 0 < ω_1_ < 1 and neutral sites with ω_1_ = 1 (Álvarez-Carretero et al., 2023). In our analysis, the ancestral branch of tetrapods was labeled as the foreground branch to test for adaptive changes potentially associated with the vertebrate transition from water to land. For the second (null) model, the same model structure is retained, but the ω_2_ parameter for the foreground branch is fixed to 1, thus not allowing positive selection. The alternative and null models were compared using a likelihood ratio test (LRT) based on their log-likelihood values. Statistical significance was assessed using the associated p-values, which were further corrected for multiple testing using the false discovery rate (FDR).

### Physiological phenotypes associated with the positively selected genes

After identifying ion channels genes showing the signature of positive selection, we sought to infer the physiological processes potentially associated with them. To this end, we used the Enrichr platform (https://maayanlab.cloud/Enrichr/) (Evangelista et al., 2023), which identifies overrepresented biological processes and pathways from a given list of gene symbols by querying approximately 240 functional annotation libraries. In this study, we compared the ion channels genes with signatures of positive selection using the MGI Mammalian Phenotype 2024 database (Baldarelli et al., 2024), which includes 11,625 genes grouped into 5,003 phenotypic categories and integrated into Enrichr. The platform applies False Discovery Rate (FDR) correction to adjust p-values, and only categories with adjusted p-values ≤ 0.01 were considered statistically significant.

### Gene family turnover rate analyses

We used the program CAFE v4.2.1 (Han et al., 2013) to identify gene family expansions and contractions in ion channels across vertebrate lineages. The program CAFE implements stochastic birth-death models of gene family evolution along a phylogeny. As input, we provided the Hierarchical Orthologous Groups (HOG) inferred with OMA, along with our time-calibrated phylogenetic tree in Newick format. Two models were evaluated: in the first, two λ (turnover rate) parameters were estimated separately for terrestrial and aquatic vertebrates. In the second, three λ turnover rate parameters were estimated, one for terrestrial vertebrates, one for aquatic vertebrates, and one for the ancestor of tetrapods. Initially, the analysis across all 86 species failed, likely due to the large number of input taxa. To resolve this issue, we reduced the dataset to 66 species while maintaining key species relevant to the water-to-land transition (Supplementary File S2). This reduced dataset allowed CAFE to successfully estimate both likelihood ratios and λ values for the gene family turnover rates across the phylogeny.

## Conflict of interest

No conflict of interest has been declared by the authors.

## Author contributions

All authors contributed equally to the conception, design, analysis, interpretation of the data, and preparation of the manuscript. All authors read and approved the final manuscript.

## Declaration of generative AI and AI-assisted technologies in the manuscript preparation process

During the preparation of this work the author(s) used ChatGPT (OpenAI) in order to improve grammar and language clarity. After using this tool, the author(s) reviewed and edited the content as needed and take(s) full responsibility for the content of the published article.

## Acknowledgments

The authors acknowledge ANID National Doctorate Scholarship 21241552 (C.U.), Fondecyt grant 1231357 (G.R.) and Fondecyt grant 1250688 (J.C.O.).

## Data Availability

Data and supplementary material are available online at https://github.com/CristobalUribe/Evolution-of-ion-channels-in-the-water-to-land-transition-of-vertebrates and https://zenodo.org/records/20075769

## Notes

https://github.com/CristobalUribe/Evolution-of-ion-channels-in-the-water-to-land-transition-of-vertebrates

https://zenodo.org/records/20075769

### Competing Interest Statement

The authors have declared no competing interest.

### Summary of Updates

Format updated: the significance of this work was added as a section.

## References

Altenhoff, A. M., Levy, J., Zarowiecki, M., Tomiczek, B., Warwick Vesztrocy, A., Dalquen, D. A., Müller, S., Telford, M. J., Glover, N. M., Dylus, D., & Dessimoz, C. (2019). OMA standalone: Orthology inference among public and custom genomes and transcriptomes. Genome Research, 29(7), 1152–1163. 10.1101/gr.243212.118

Álvarez-Carretero, S., Kapli, P., & Yang, Z. (2023). Beginner’s Guide on the Use of PAML to Detect Positive Selection. Molecular Biology and Evolution, 40(4), msad041. 10.1093/molbev/msad041

Amburgey, K., Bailey, A., Hwang, J. H., Tarnopolsky, M. A., Bonnemann, C. G., Medne, L., Mathews, K. D., Collins, J., Daube, J. R., Wellman, G. P., Callaghan, B., Clarke, N. F., & Dowling, J. J. (2013). Genotype-phenotype correlations in recessive RYR1-related myopathies. Orphanet Journal of Rare Diseases, 8(1), 117. 10.1186/1750-1172-8-117

Amores, A., Force, A., Yan, Y.-L., Joly, L., Amemiya, C., Fritz, A., Ho, R. K., Langeland, J., Prince, V., Wang, Y.-L., Westerfield, M., Ekker, M., & Postlethwait, J. H. (1998). Zebrafish *hox* Clusters and Vertebrate Genome Evolution. Science, 282(5394), 1711–1714. 10.1126/science.282.5394.1711

Aristide, L., & Fernández, R. (2023). Genomic Insights into Mollusk Terrestrialization: Parallel and Convergent Gene Family Expansions as Key Facilitators in Out-of-the-Sea Transitions. Genome Biology and Evolution, 15(10), evad176. 10.1093/gbe/evad176

Ashley-Ross, M. A., Hsieh, S. T., Gibb, A. C., & Blob, R. W. (2013a). Vertebrate Land Invasions-Past, Present, and Future: An Introduction to the Symposium. Integrative and Comparative Biology, 53(2), 192–196. 10.1093/icb/ict048

Ashley-Ross, M. A., Hsieh, S. T., Gibb, A. C., & Blob, R. W. (2013b). Vertebrate Land Invasions-Past, Present, and Future: An Introduction to the Symposium. Integrative and Comparative Biology, 53(2), 192–196. 10.1093/icb/ict048

Baldarelli, R. M., Smith, C. L., Ringwald, M., Richardson, J. E., Bult, C. J., Mouse Genome Informatics Group, Anagnostopoulos, A., Begley, D. A., Bello, S. M., Christie, K., Finger, J. H., Hale, P., Hayamizu, T. F., Hill, D. P., Knowlton, M. N., Krupke, D. M., McAndrews, M., Law, M., McCright, I. J., … Ormsby, J. E. (2024). Mouse Genome Informatics: An integrated knowledgebase system for the laboratory mouse. GENETICS, 227(1), iyae031. 10.1093/genetics/iyae031

Barker, B. S., Young, G. T., Soubrane, C. H., Stephens, G. J., Stevens, E. B., & Patel, M. K. (2017). Ion Channels. En Conn’s Translational Neuroscience (pp. 11–43). Elsevier. 10.1016/B978-0-12-802381-5.00002-6

Barnes, N. M., Hales, T. G., Lummis, S. C. R., & Peters, J. A. (2009). The 5-HT3 receptor – the relationship between structure and function. Neuropharmacology, 56(1), 273–284. 10.1016/j.neuropharm.2008.08.003

Bhanuprakash, V., Chhotaray, S., Pruthviraj, D. R., Rawat, C., Karthikeyan, A., & Panigrahi, M. (2018). Copy number variation in livestock: A mini review. Veterinary World, 11(4), 535–541. 10.14202/vetworld.2018.535-541

Bhattacharya, A., Dang, H., Zhu, Q.-M., Schnegelsberg, B., Rozengurt, N., Cain, G., Prantil, R., Vorp, D. A., Guy, N., Julius, D., Ford, A. P. D. W., Lester, H. A., & Cockayne, D. A. (2004). Uropathic observations in mice expressing a constitutively active point mutation in the 5-HT3A receptor subunit. The Journal of Neuroscience: The Official Journal of the Society for Neuroscience, 24(24), 5537–5548. 10.1523/JNEUROSCI.5658-03.2004

Bondareva, O., Petrova, T., Bodrov, S., Gavrilo, M., Smorkatcheva, A., & Abramson, N. (2023). How voles adapt to subterranean lifestyle: Insights from RNA-seq. Frontiers in Ecology and Evolution, 11, 1085993. 10.3389/fevo.2023.1085993

Boudewyn, L. C., Sikora, J., Kuchar, L., Ledvinova, J., Grishchuk, Y., Wang, S. L., Dobrenis, K., & Walkley, S. U. (2017). N-butyldeoxynojirimycin delays motor deficits, cerebellar microgliosis, and Purkinje cell loss in a mouse model of mucolipidosis type IV. Neurobiology of Disease, 105, 257–270. 10.1016/j.nbd.2017.06.003

Brun, L., Viemari, J.-C., & Villard, L. (2022). Mouse models of Kcnq2 dysfunction. Epilepsia, 63(11), 2813–2826. 10.1111/epi.17405

Cabezas-Bratesco, D., Mcgee, F. A., Colenso, C. K., Zavala, K., Granata, D., Carnevale, V., Opazo, J. C., & Brauchi, S. E. (2022). Sequence and structural conservation reveal fingerprint residues in TRP channels. eLife, 11, e73645. 10.7554/eLife.73645

Cancino, S., Cabeza-Bratesco, D., Matamala, E., Jork, J. M., Zavala, K., Montecinos, C., Bustamante-Villarreal, S., Maldonado, R., Riadi, G., Brauchi, S. E., & Opazo, J. C. (2025). An Evolutionary Novelty in TRPV1 Functional Regulation: Characterization of a Dominant-Negative Isoform Exclusive to Catarrhine Primates. Evolutionary Biology. 10.1101/2025.01.24.634766

Castillo, K., Diaz-Franulic, I., Canan, J., Gonzalez-Nilo, F., & Latorre, R. (2018). Thermally activated TRP channels: Molecular sensors for temperature detection. Physical Biology, 15(2), 021001. 10.1088/1478-3975/aa9a6f

Catacuzzeno, L., Michelucci, A., & Franciolini, F. (2025). The crucial decade that ion channels were proven to exist: The vision of Bertil Hille and Clay Armstrong and how it came through. Pflügers Archiv - European Journal of Physiology, 477(7), 903–917. 10.1007/s00424-025-03085-5

Chagovetz, A. A., Klatt Shaw, D., Ritchie, E., Hoshijima, K., & Grunwald, D. J. (2019). Interactions among ryanodine receptor isotypes contribute to muscle fiber type development and function. Disease Models & Mechanisms, 13(2), dmm038844. 10.1242/dmm.038844

Chen, D. Y., Chowdhury, S., Farnaes, L., Friedman, J. R., Honold, J., Dimmock, D. P., & Gold, O. B. O. T. R. I. J. J. (2018). Rapid Diagnosis of KCNQ2-Associated Early Infantile Epileptic Encephalopathy Improved Outcome. Pediatric Neurology, 86, 69–70. 10.1016/j.pediatrneurol.2018.06.002

Chen, L., DeVries, A. L., & Cheng, C. H. (1997). Evolution of antifreeze glycoprotein gene from a trypsinogen gene in Antarctic notothenioid fish. Proceedings of the National Academy of Sciences of the United States of America, 94(8), 3811–3816. 10.1073/pnas.94.8.3811

Chen, Y.-C., Kung, S.-S., Chen, B.-Y., Hung, C.-C., Chen, C.-C., Wang, T.-Y., Wu, Y.-M., Lin, W.- H., Tzeng, C.-S., & Chow, W.-Y. (2001). Identifications, Classification, and Evolution of the Vertebrate α-Amino-3-Hydroxy-5-Methyl-4-Isoxazole Propionic Acid (AMPA) Receptor Subunit Genes. Journal of Molecular Evolution, 53(6), 690–702. 10.1007/s002390010256

Clack, J. A. (2006). The emergence of early tetrapods. Palaeogeography, Palaeoclimatology, Palaeoecology, 232(2-4), 167–189. 10.1016/j.palaeo.2005.07.019

Clack, J. A. (2009). The Fish–Tetrapod Transition: New Fossils and Interpretations. Evolution: Education and Outreach, 2(2), 213–223. 10.1007/s12052-009-0119-2

Corliss, B. A., Delalio, L. J., Stevenson Keller, T. C., Keller, A. S., Keller, D. A., Corliss, B. H., Beers, J. M., Peirce, S. M., & Isakson, B. E. (2019). Vascular Expression of Hemoglobin Alpha in Antarctic Icefish Supports Iron Limitation as Novel Evolutionary Driver. Frontiers in Physiology, 10, 1389. 10.3389/fphys.2019.01389

Csekő, K., Beckers, B., Keszthelyi, D., & Helyes, Z. (2019). Role of TRPV1 and TRPA1 Ion Channels in Inflammatory Bowel Diseases: Potential Therapeutic Targets? *Pharmaceuticals (Basel*, Switzerland), 12(2), 48. 10.3390/ph12020048

Cupello, C., Hirasawa, T., Tatsumi, N., Yabumoto, Y., Gueriau, P., Isogai, S., Matsumoto, R., Saruwatari, T., King, A., Hoshino, M., Uesugi, K., Okabe, M., & Brito, P. M. (2022). Lung evolution in vertebrates and the water-to-land transition. eLife, 11, e77156. 10.7554/eLife.77156

Delmas, P., & Coste, B. (2013). Mechano-Gated Ion Channels in Sensory Systems. Cell, 155(2), 278–284. 10.1016/j.cell.2013.09.026

Devaux, J. J., Kleopa, K. A., Cooper, E. C., & Scherer, S. S. (2004). KCNQ2 is a nodal K+ channel. The Journal of Neuroscience: The Official Journal of the Society for Neuroscience, 24(5), 1236–1244. 10.1523/JNEUROSCI.4512-03.2004

Du, K., Yang, L., & He, S. (2015). Phylogenomic analyses reveal a molecular signature linked to subterranean adaptation in rodents. BMC Evolutionary Biology, 15(1), 287. 10.1186/s12862-015-0564-1

Dubey, N. K., Kumar, V., & Goswami, C. (2024). Sperm-Specific CatSper is Not Conserved in All Vertebrates and May Not be the Only Progesterone-Responsive Ion Channel Present in Sperm. The Journal of Membrane Biology, 257(3-4), 215–230. 10.1007/s00232-024-00316-1

Evangelista, J. E., Xie, Z., Marino, G. B., Nguyen, N., Clarke, D. J. B., & Ma’ayan, A. (2023). Enrichr-KG: Bridging enrichment analysis across multiple libraries. Nucleic Acids Research, 51(W1), W168–W179. 10.1093/nar/gkad393

Flores-Aldama, L., Vandewege, M. W., Zavala, K., Colenso, C. K., Gonzalez, W., Brauchi, S. E., & Opazo, J. C. (2020). Evolutionary analyses reveal independent origins of gene repertoires and structural motifs associated to fast inactivation in calcium-selective TRPV channels. Scientific Reports, 10(1), 8684. 10.1038/s41598-020-65679-6

Gouin, O., L’Herondelle, K., Lebonvallet, N., Le Gall-Ianotto, C., Sakka, M., Buhé, V., Plée-Gautier, E., Carré, J.-L., Lefeuvre, L., Misery, L., & Le Garrec, R. (2017). TRPV1 and TRPA1 in cutaneous neurogenic and chronic inflammation: Pro-inflammatory response induced by their activation and their sensitization. Protein & Cell, 8(9), 644–661. 10.1007/s13238-017-0395-5

Gregory, T. R. (2001). Coincidence, coevolution, or causation? DNA content, cellsize, and the C-value enigma. Biological Reviews, 76(1), 65–101. 10.1111/j.1469-185X.2000.tb00059.x

Gregory, T. R. (2002). A BIRD’S-EYE VIEW OF THE C-VALUE ENIGMA: GENOME SIZE, CELL SIZE, AND METABOLIC RATE IN THE CLASS AVES. Evolution, 56(1), 121–130. 10.1111/j.0014-3820.2002.tb00854.x

Hamed, Y. M. F., Ghosh, B., & Marshall, K. L. (2024). PIEZO ion channels: Force sensors of the interoceptive nervous system. The Journal of Physiology, 602(19), 4777–4788. 10.1113/JP284077

Han, M. V., Thomas, G. W. C., Lugo-Martinez, J., & Hahn, M. W. (2013). Estimating gene gain and loss rates in the presence of error in genome assembly and annotation using CAFE 3. Molecular Biology and Evolution, 30(8), 1987–1997. 10.1093/molbev/mst100

Harrison, P. W., Amode, M. R., Austine-Orimoloye, O., Azov, A. G., Barba, M., Barnes, I., Becker, A., Bennett, R., Berry, A., Bhai, J., Bhurji, S. K., Boddu, S., Branco Lins, P. R., Brooks, L., Ramaraju, S. B., Campbell, L. I., Martinez, M. C., Charkhchi, M., Chougule, K., … Yates, A. D. (2024). Ensembl 2024. Nucleic Acids Research, 52(D1), D891–D899. 10.1093/nar/gkad1049

Hartmann, B., Ahmadi, S., Heppenstall, P. A., Lewin, G. R., Schott, C., Borchardt, T., Seeburg, P. H., Zeilhofer, H. U., Sprengel, R., & Kuner, R. (2004). The AMPA Receptor Subunits GluR-A and GluR-B Reciprocally Modulate Spinal Synaptic Plasticity and Inflammatory Pain. Neuron, 44(4), 637–650. 10.1016/j.neuron.2004.10.029

Hoegg, S., & Meyer, A. (2007). Phylogenomic analyses of KCNA gene clusters in vertebrates: Why do gene clusters stay intact? BMC Evolutionary Biology, 7(1), 139. 10.1186/1471-2148-7-139

Huang, S. M., Lee, H., Chung, M.-K., Park, U., Yu, Y. Y., Bradshaw, H. B., Coulombe, P. A., Walker, J. M., & Caterina, M. J. (2008). Overexpressed transient receptor potential vanilloid 3 ion channels in skin keratinocytes modulate pain sensitivity via prostaglandin E2. The Journal of Neuroscience: The Official Journal of the Society for Neuroscience, 28(51), 13727–13737. 10.1523/JNEUROSCI.5741-07.2008

Hughes, A. L., & Friedman, R. (2008). Genome Size Reduction in the Chicken Has Involved Massive Loss of Ancestral Protein-Coding Genes. Molecular Biology and Evolution, 25(12), 2681–2688. 10.1093/molbev/msn207

Jaillon, O., Aury, J.-M., Brunet, F., Petit, J.-L., Stange-Thomann, N., Mauceli, E., Bouneau, L., Fischer, C., Ozouf-Costaz, C., Bernot, A., Nicaud, S., Jaffe, D., Fisher, S., Lutfalla, G., Dossat, C., Segurens, B., Dasilva, C., Salanoubat, M., Levy, M., … Roest Crollius, H. (2004). Genome duplication in the teleost fish Tetraodon nigroviridis reveals the early vertebrate proto-karyotype. Nature, 431(7011), 946–957. 10.1038/nature03025

Jiang, M., Shi, L., Li, X., Dong, Q., Sun, H., Du, Y., Zhang, Y., Shao, T., Cheng, H., Chen, W., & Wang, Z. (2020). Genome-wide adaptive evolution to underground stresses in subterranean mammals: Hypoxia adaption, immunity promotion, and sensory specialization. Ecology and Evolution, 10(14), 7377–7388. 10.1002/ece3.6462

Kalinovskii, A. P., Utkina, L. L., Korolkova, Y. V., & Andreev, Y. A. (2023). TRPV3 Ion Channel: From Gene to Pharmacology. International Journal of Molecular Sciences, 24(10), 8601. 10.3390/ijms24108601

Kashio, M., & Tominaga, M. (2022). TRP channels in thermosensation. Current Opinion in Neurobiology, 75, 102591. 10.1016/j.conb.2022.102591

Katoh, K., & Standley, D. M. (2013). MAFFT multiple sequence alignment software version 7: Improvements in performance and usability. Molecular Biology and Evolution, 30(4), 772–780. 10.1093/molbev/mst010

Kauer, J. A., & Gibson, H. E. (2009). Hot flash: TRPV channels in the brain. Trends in Neurosciences, 32(4), 215–224. 10.1016/j.tins.2008.12.006

Kim, Y.-J., Choi, S. J., Hong, S.-I., Park, J.-C., Lee, Y., Ma, S.-X., Hur, K.-H., Lee, Y., Kim, K.-M., Kim, H. K., Kim, H. Y., Lee, S.-Y., Choi, S.-Y., & Jang, C.-G. (2024). The ion channel TRPA1 is a modulator of the cocaine reward circuit in the nucleus accumbens. Molecular Psychiatry, 29(11), 3607–3622. 10.1038/s41380-024-02623-4

Kobayashi, T., Tsutsumi, A., Kurebayashi, N., Saito, K., Kodama, M., Sakurai, T., Kikkawa, M., Murayama, T., & Ogawa, H. (2022). Molecular basis for gating of cardiac ryanodine receptor explains the mechanisms for gain- and loss-of function mutations. Nature Communications, 13(1), 2821. 10.1038/s41467-022-30429-x

Krogh, A., Larsson, B., Von Heijne, G., & Sonnhammer, E. L. L. (2001). Predicting transmembrane protein topology with a hidden markov model: Application to complete genomes11Edited by F. Cohen. Journal of Molecular Biology, 305(3), 567–580. 10.1006/jmbi.2000.4315

Kumar, S., Suleski, M., Craig, J. M., Kasprowicz, A. E., Sanderford, M., Li, M., Stecher, G., & Hedges, S. B. (2022). TimeTree 5: An Expanded Resource for Species Divergence Times. Molecular Biology and Evolution, 39(8), msac174. 10.1093/molbev/msac174

Kuraku, S., & Meyer, A. (2009). The evolution and maintenance of Hox gene clusters in vertebrates and the teleost-specific genome duplication. The International Journal of Developmental Biology, 53(5-6), 765–773. 10.1387/ijdb.072533km

Lanner, J. T., Georgiou, D. K., Joshi, A. D., & Hamilton, S. L. (2010). Ryanodine Receptors: Structure, Expression, Molecular Details, and Function in Calcium Release. Cold Spring Harbor Perspectives in Biology, 2(11), a003996–a003996. 10.1101/cshperspect.a003996

Laursen, W. J., Schneider, E. R., Merriman, D. K., Bagriantsev, S. N., & Gracheva, E. O. (2016). Low-cost functional plasticity of TRPV1 supports heat tolerance in squirrels and camels. Proceedings of the National Academy of Sciences, 113(40), 11342–11347. 10.1073/pnas.1604269113

Lee, K., Jo, Y. Y., Chung, G., Jung, J. H., Kim, Y. H., & Park, C.-K. (2021). Functional Importance of Transient Receptor Potential (TRP) Channels in Neurological Disorders. Frontiers in Cell and Developmental Biology, 9, 611773. 10.3389/fcell.2021.611773

Lei, J., & Tominaga, M. (2025). TRPV3 in skin thermosensation and temperature responses. The Journal of Physiological Sciences, 75(1), 100005. 10.1016/j.jphyss.2025.100005

Lovell, P. V., Wirthlin, M., Wilhelm, L., Minx, P., Lazar, N. H., Carbone, L., Warren, W. C., & Mello, C. V. (2014). Conserved syntenic clusters of protein coding genes are missing in birds. Genome Biology, 15(12), 565. 10.1186/s13059-014-0565-1

Lu, X., Yao, Z., Wang, Y., Yin, C., Li, J., Chai, L., Dong, W., Yuan, L., Lai, R., & Yang, S. (2022). The acquisition of cold sensitivity during TRPM8 ion channel evolution. Proceedings of the National Academy of Sciences, 119(21), e2201349119. 10.1073/pnas.2201349119

Luo, L., Wang, Y., Li, B., Xu, L., Kamau, P. M., Zheng, J., Yang, F., Yang, S., & Lai, R. (2019). Molecular basis for heat desensitization of TRPV1 ion channels. Nature Communications, 10(1). 10.1038/s41467-019-09965-6

Ma, D., Zheng, Y., Li, X., Zhou, X., Yang, Z., Zhang, Y., Wang, L., Zhang, W., Fang, J., Zhao, G., Hou, P., Nan, F., Yang, W., Su, N., Gao, Z., & Guo, J. (2023). Ligand activation mechanisms of human KCNQ2 channel. Nature Communications, 14(1), 6632. 10.1038/s41467-023-42416-x

MacIver, M. A., & Finlay, B. L. (2022). The neuroecology of the water-to-land transition and the evolution of the vertebrate brain. Philosophical Transactions of the Royal Society B: Biological Sciences, 377(1844), 20200523. 10.1098/rstb.2020.0523

Mandadi, S., Sokabe, T., Shibasaki, K., Katanosaka, K., Mizuno, A., Moqrich, A., Patapoutian, A., Fukumi-Tominaga, T., Mizumura, K., & Tominaga, M. (2009). TRPV3 in keratinocytes transmits temperature information to sensory neurons via ATP. Pflügers Archiv - European Journal of Physiology, 458(6), 1093–1102. 10.1007/s00424-009-0703-x

Mao, M., Jancovski, N., Kushner, Y., Teasdale, L., Truong, P., Zhou, K., Reid, S., Jia, L., Aung, Y. H., Li, M., Reid, C. A., Byars, S., Scheffer, I., Petrou, S., & Maljevic, S. (2025). Developmental dysfunction in a preclinical model of Kcnq2 developmental and epileptic encephalopathy. Neurobiology of Disease, 205, 106782. 10.1016/j.nbd.2024.106782

Marban, E., Yamagishi, T., & Tomaselli, G. F. (1998). Structure and function of voltage-gated sodium channels. The Journal of Physiology, 508 (Pt 3)(Pt 3), 647-657. 10.1111/j.1469-7793.1998.647bp.x

Marchler-Bauer, A., Lu, S., Anderson, J. B., Chitsaz, F., Derbyshire, M. K., DeWeese-Scott, C., Fong, J. H., Geer, L. Y., Geer, R. C., Gonzales, N. R., Gwadz, M., Hurwitz, D. I., Jackson, J. D., Ke, Z., Lanczycki, C. J., Lu, F., Marchler, G. H., Mullokandov, M., Omelchenko, M. V., … Bryant, S. H. (2011). CDD: A Conserved Domain Database for the functional annotation of proteins. Nucleic Acids Research, 39(Database), D225–D229. 10.1093/nar/gkq1189

Martínez-Redondo, G. I., Simón Guerrero, C., Aristide, L., Balart-García, P., Tonzo, V., & Fernández, R. (2023). Parallel duplication and loss of aquaporin-coding genes during the “out of the sea” transition as potential key drivers of animal terrestrialization. Molecular Ecology, 32(8), 2022–2040. 10.1111/mec.16854

McGowen, M. R., Grossman, L. I., & Wildman, D. E. (2012). Dolphin genome provides evidence for adaptive evolution of nervous system genes and a molecular rate slowdown. Proceedings of the Royal Society B: Biological Sciences, 279(1743), 3643–3651. 10.1098/rspb.2012.0869

Medeiros-Domingo, A., Tan, B.-H., Crotti, L., Tester, D. J., Eckhardt, L., Cuoretti, A., Kroboth, S. L., Song, C., Zhou, Q., Kopp, D., Schwartz, P. J., Makielski, J. C., & Ackerman, M. J. (2010). Gain-of-function mutation S422L in the KCNJ8-encoded cardiac KATP channel Kir6.1 as a pathogenic substrate for J-wave syndromes. Heart Rhythm, 7(10), 1466–1471. 10.1016/j.hrthm.2010.06.016

Nelson, P. L., Beck, A., & Cheng, H. (2011). Transient receptor proteins illuminated: Current views on TRPs and disease. The Veterinary Journal, 187(2), 153–164. 10.1016/j.tvjl.2010.01.020

Nery, M. F., González, D. J., & Opazo, J. C. (2013). How to Make a Dolphin: Molecular Signature of Positive Selection in Cetacean Genome. PLoS ONE, 8(6), e65491. 10.1371/journal.pone.0065491

Nichols, C. G. (2016). Adenosine Triphosphate-Sensitive Potassium Currents in Heart Disease and Cardioprotection. Cardiac Electrophysiology Clinics, 8(2), 323–335. 10.1016/j.ccep.2016.01.005

O’Connor, T. N., Van Den Bersselaar, L. R., Chen, Y. S., Nicolau, S., Simon, B., Huseth, A., Todd, J. J., Van Petegem, F., Sarkozy, A., Goldberg, M. F., Voermans, N. C., Dirksen, R. T., & RYR1 Myopathy Consortium Leslie Biesecker Bethesda, US, Johann Böhm Strasbourg, FR, Carsten Bönnemann Bethesda, US, Oliver Clarke New York, US, Razvan Cornea Minneapolis, US, Robert Dirksen Rochester, US, James Dowling Toronto, CA, Michael Goldberg Pittsburgh, US, Susan Hamilton Houston, US, Drew Huseth Mesa, US, Heinz Jungbluth London, UK, Tokunbor Lawal Bethesda, US, Andrew Marks New York, US, Isabelle Marty Grenoble, FR, Livija Medne Philadelphia, US, Eva Michael Gothenburg, SE, Payam Mohassel Bethesda,. (2023). RYR-1-Related Diseases International Research Workshop: From Mechanisms to Treatments Pittsburgh, PA, U.S.A., 21-22 July 2022. Journal of Neuromuscular Diseases, 10(1), 135–154. 10.3233/JND-221609

Pajic, P., Pavlidis, P., Dean, K., Neznanova, L., Romano, R.-A., Garneau, D., Daugherity, E., Globig, A., Ruhl, S., & Gokcumen, O. (2019). Independent amylase gene copy number bursts correlate with dietary preferences in mammals. eLife, 8, e44628. 10.7554/eLife.44628

Palchevskyi, S., Czarnocki-Cieciura, M., Vistoli, G., Gervasoni, S., Nowak, E., Beccari, A. R., Nowotny, M., & Talarico, C. (2023). Structure of human TRPM8 channel. Communications Biology, 6(1). 10.1038/s42003-023-05425-6

Patapoutian, A., Peier, A. M., Story, G. M., & Viswanath, V. (2003). ThermoTRP channels and beyond: Mechanisms of temperature sensation. Nature Reviews Neuroscience, 4(7), 529–539. 10.1038/nrn1141

Perry, G. H., Dominy, N. J., Claw, K. G., Lee, A. S., Fiegler, H., Redon, R., Werner, J., Villanea, F. A., Mountain, J. L., Misra, R., Carter, N. P., Lee, C., & Stone, A. C. (2007). Diet and the evolution of human amylase gene copy number variation. Nature Genetics, 39(10), 1256–1260. 10.1038/ng2123

Pifferi, S., Boccaccio, A., & Menini, A. (2006). Cyclic nucleotide-gated ion channels in sensory transduction. FEBS Letters, 580(12), 2853–2859. 10.1016/j.febslet.2006.03.086

Popova, E. (2014). Ionotropic GABA Receptors and Distal Retinal ON and OFF Responses. Scientifica, 2014, 149187. 10.1155/2014/149187

Premkumar, L. S. (2014). Transient Receptor Potential Channels as Targets for Phytochemicals. ACS Chemical Neuroscience, 5(11), 1117–1130. 10.1021/cn500094a

Ribelayga, C. (2010). Vertebrate Vision: TRP Channels in the Spotlight. Current Biology, 20(6), R278–R280. 10.1016/j.cub.2010.02.012

Rocha, J. L., Godinho, R., Brito, J. C., & Nielsen, R. (2021). Life in Deserts: The Genetic Basis of Mammalian Desert Adaptation. Trends in Ecology & Evolution, 36(7), 637–650. 10.1016/j.tree.2021.03.007

Rossetti, S., Chauveau, D., Kubly, V., Slezak, J. M., Saggar-Malik, A. K., Pei, Y., Ong, A. C., Stewart, F., Watson, M. L., Bergstralh, E. J., Winearls, C. G., Torres, V. E., & Harris, P. C. (2003). Association of mutation position in polycystic kidney disease 1 (PKD1) gene and development of a vascular phenotype. The Lancet, 361(9376), 2196–2201. 10.1016/S0140-6736(03)13773-7

Saito, S., Fukuta, N., Shingai, R., & Tominaga, M. (2011). Evolution of Vertebrate Transient Receptor Potential Vanilloid 3 Channels: Opposite Temperature Sensitivity between Mammals and Western Clawed Frogs. PLoS Genetics, 7(4), e1002041. 10.1371/journal.pgen.1002041

Saito, S., Ohkita, M., Saito, C. T., Takahashi, K., Tominaga, M., & Ohta, T. (2016). Evolution of Heat Sensors Drove Shifts in Thermosensation between Xenopus Species Adapted to Different Thermal Niches. Journal of Biological Chemistry, 291(21), 11446–11459. 10.1074/jbc.m115.702498

Saito, S., & Saito, C. T. (2025). Evolution of Temperature Receptors and Their Roles in Sensory Diversification and Adaptation. Zoological Science, 42(1). 10.2108/zs240060

Saito, S., Saito, C. T., Nozawa, M., & Tominaga, M. (2019). Elucidating the functional evolution of heat sensors among*Xenopus*species adapted to different thermal niches by ancestral sequence reconstruction. Molecular Ecology, 28(15), 3561–3571. 10.1111/mec.15170

Saito, S., & Tominaga, M. (2017). Evolutionary tuning of TRPA1 and TRPV1 thermal and chemical sensitivity in vertebrates. Temperature, 4(2), 141–152. 10.1080/23328940.2017.1315478

Sato, T., Arimura, T., Murata, K., Kawamura, M., Obama, W., Suzuki, M., Nakauchi, Y., Tominaga, A., Morita, M., Hiraoka, K., Takayama-Watanabe, E., & Watanabe, A. (2021). Differences of Extracellular Cues and Ca2+ Permeable Channels in the Signaling Pathways for Inducing Amphibian Sperm Motility. Zoological Science, 38(4). 10.2108/zs200159

Sayers, E. W., Beck, J., Bolton, E. E., Brister, J. R., Chan, J., Connor, R., Feldgarden, M., Fine, A. M., Funk, K., Hoffman, J., Kannan, S., Kelly, C., Klimke, W., Kim, S., Lathrop, S., Marchler-Bauer, A., Murphy, T. D., O’Sullivan, C., Schmieder, E., … Pruitt, K. D. (2025). Database resources of the National Center for Biotechnology Information in 2025. Nucleic Acids Research, 53(D1), D20–D29. 10.1093/nar/gkae979

Scicchitano, P., Carbonara, S., Ricci, G., Mandurino, C., Locorotondo, M., Bulzis, G., Gesualdo, M., Zito, A., Carbonara, R., Dentamaro, I., Riccioni, G., & Ciccone, M. M. (2012). HCN channels and heart rate. *Molecules (Basel*, Switzerland), 17(4), 4225–4235. 10.3390/molecules17044225

Shin, S. C., Kim, S., Kim, H.-W., Lee, J. H., & Kim, J.-H. (2024). Gene loss in Antarctic icefish: Evolutionary adaptations mimicking Fanconi Anemia? BMC Genomics, 25(1), 1102. 10.1186/s12864-024-11028-0

Shukla, K. K., Mahdi, A. A., & Rajender, S. (2012). Ion Channels in Sperm Physiology and Male Fertility and Infertility. Journal of Andrology, 33(5), 777–788. 10.2164/jandrol.111.015552

Silverman, H. A., Chen, A., Kravatz, N. L., Chavan, S. S., & Chang, E. H. (2020). Involvement of Neural Transient Receptor Potential Channels in Peripheral Inflammation. Frontiers in Immunology, 11, 590261. 10.3389/fimmu.2020.590261

Stelzer, G., Rosen, N., Plaschkes, I., Zimmerman, S., Twik, M., Fishilevich, S., Stein, T. I., Nudel, R., Lieder, I., Mazor, Y., Kaplan, S., Dahary, D., Warshawsky, D., Guan-Golan, Y., Kohn, A., Rappaport, N., Safran, M., & Lancet, D. (2016). The GeneCards Suite: From Gene Data Mining to Disease Genome Sequence Analyses. Current Protocols in Bioinformatics, 54, 1.30.1-1.30.33. 10.1002/cpbi.5

Straub, R. H. (2014). TRPV1, TRPA1, and TRPM8 channels in inflammation, energy redirection, and water retention: Role in chronic inflammatory diseases with an evolutionary perspective. Journal of Molecular Medicine, 92(9), 925-937. 10.1007/s00109-014-1175-9

Sun, Y.-B., Zhou, W.-P., Liu, H.-Q., Irwin, D. M., Shen, Y.-Y., & Zhang, Y.-P. (2013). Genome-Wide Scans for Candidate Genes Involved in the Aquatic Adaptation of Dolphins. Genome Biology and Evolution, 5(1), 130–139. 10.1093/gbe/evs123

Suyama, M., Torrents, D., & Bork, P. (2006). PAL2NAL: Robust conversion of protein sequence alignments into the corresponding codon alignments. Nucleic Acids Research, 34(Web Server issue), W609-612. 10.1093/nar/gkl315

Syrbe, S., Hedrich, U. B. S., Riesch, E., Djémié, T., Müller, S., Møller, R. S., Maher, B., Hernandez-Hernandez, L., Synofzik, M., Caglayan, H. S., Arslan, M., Serratosa, J. M., Nothnagel, M., May, P., Krause, R., Löffler, H., Detert, K., Dorn, T., Vogt, H., … Lemke, J. R. (2015). De novo loss- or gain-of-function mutations in KCNA2 cause epileptic encephalopathy. Nature Genetics, 47(4), 393–399. 10.1038/ng.3239

Tester, D. J., Tan, B.-H., Medeiros-Domingo, A., Song, C., Makielski, J. C., & Ackerman, M. J. (2011). Loss-of-Function Mutations in the *KCNJ8* -Encoded Kir6.1 K_ATP_ Channel and Sudden Infant Death Syndrome. Circulation: Cardiovascular Genetics, 4(5), 510–515. 10.1161/CIRCGENETICS.111.960195

Thompson, A. J., & Lummis, S. C. R. (2006). 5-HT3 receptors. Current Pharmaceutical Design, 12(28), 3615–3630. 10.2174/138161206778522029

Tsagkogeorga, G., McGowen, M. R., Davies, K. T. J., Jarman, S., Polanowski, A., Bertelsen, M. F., & Rossiter, S. J. (2015). A phylogenomic analysis of the role and timing of molecular adaptation in the aquatic transition of cetartiodactyl mammals. Royal Society Open Science, 2(9), 150156. 10.1098/rsos.150156

Uribe, C., Nery, M. F., Zavala, K., Mardones, G. A., Riadi, G., & Opazo, J. C. (2024). Evolution of ion channels in cetaceans: A natural experiment in the tree of life. Scientific Reports, 14(1), 17024. 10.1038/s41598-024-66082-1

Vay, L., Gu, C., & McNaughton, P. A. (2012). The thermo-TRP ion channel family: Properties and therapeutic implications. British Journal of Pharmacology, 165(4), 787–801. 10.1111/j.1476-5381.2011.01601.x

Vicente-Carrillo, A., Álvarez-Rodríguez, M., & Rodriguez-Martinez, H. (2023). The Cation/Calcium Channel of Sperm (CatSper): A Common Role Played Despite Inter-Species Variation? International Journal of Molecular Sciences, 24(18), 13750. 10.3390/ijms241813750

Vornanen, M. (2006). Temperature and Ca^2+^ dependence of ^[3^ H]ryanodine binding in the burbot (*Lota lota* L.) heart. *American Journal of Physiology-Regulatory*, Integrative and Comparative Physiology, 290(2), R345–R351. 10.1152/ajpregu.00443.2005

Wahl-Schott, C., & Biel, M. (2009). HCN channels: Structure, cellular regulation and physiological function. Cellular and Molecular Life Sciences, 66(3), 470–494. 10.1007/s00018-008-8525-0

Wang, H., McGoldrick, L. L., & Chung, J.-J. (2021). Sperm ion channels and transporters in male fertility and infertility. Nature Reviews. Urology, 18(1), 46–66. 10.1038/s41585-020-00390-9

Wang, J., Chitsaz, F., Derbyshire, M. K., Gonzales, N. R., Gwadz, M., Lu, S., Marchler, G. H., Song, J. S., Thanki, N., Yamashita, R. A., Yang, M., Zhang, D., Zheng, C., Lanczycki, C. J., & Marchler-Bauer, A. (2023). The conserved domain database in 2023. Nucleic Acids Research, 51(D1), D384–D388. 10.1093/nar/gkac1096

Wang, X., Wang, Y., Xu, T., Fan, Y., Ding, Y., & Qian, J. (2023). A novel compound heterozygous mutation of the CLCN7 gene is associated with autosomal recessive osteopetrosis. Frontiers in Pediatrics, 11, 978879. 10.3389/fped.2023.978879

Willis, W. D. (2009). The role of TRPV1 receptors in pain evoked by noxious thermal and chemical stimuli. Experimental Brain Research, 196(1), 5–11. 10.1007/s00221-009-1760-2

Wu, C.-K., Lin, J.-F., Lee, T.-S., Kou, Y. R., & Tarng, D.-C. (2021). Role of TRPA1 in Tissue Damage and Kidney Disease. International Journal of Molecular Sciences, 22(7), 3415. 10.3390/ijms22073415

Wu, L., Jiao, X., Zhang, D., Cheng, Y., Song, G., Qu, Y., & Lei, F. (2021). Comparative Genomics and Evolution of Avian Specialized Traits. Current Genomics, 22(7), 496–511. 10.2174/1389202923666211227143952

Xie, G., Harrison, J., Clapcote, S. J., Huang, Y., Zhang, J.-Y., Wang, L.-Y., & Roder, J. C. (2010). A New Kv1.2 Channelopathy Underlying Cerebellar Ataxia. Journal of Biological Chemistry, 285(42), 32160–32173. 10.1074/jbc.M110.153676

Yang, H.-T., Tweedie, D., Wang, S., Guia, A., Vinogradova, T., Bogdanov, K., Allen, P. D., Stern, M. D., Lakatta, E. G., & Boheler, K. R. (2002). The ryanodine receptor modulates the spontaneous beating rate of cardiomyocytes during development. Proceedings of the National Academy of Sciences of the United States of America, 99(14), 9225–9230. 10.1073/pnas.142651999

Yang, Z. (2007). PAML 4: Phylogenetic Analysis by Maximum Likelihood. Molecular Biology and Evolution, 24(8), 1586–1591. 10.1093/molbev/msm088

Ye, J., Tang, S., Miao, P., Gong, Z., Shu, Q., Feng, J., & Li, Y. (2023). Clinical analysis and functional characterization of KCNQ2-related developmental and epileptic encephalopathy. Frontiers in Molecular Neuroscience, 16, 1205265. 10.3389/fnmol.2023.1205265

Yoshida, M., & Yoshida, K. (2025). Activation of motility and chemotaxis in the spermatozoa. Reproductive Medicine and Biology, 24(1), e12638. 10.1002/rmb2.12638

Yuan, S., Zhang, R., Jin, Y., Li, X., Li, L., Zhang, D., Ling, Y., Zhang, K., Wu, X., Zhang, X., & Fu, H. (2025). Multiomic Analyses Reveal the Molecular Mechanisms of Arid Adaptation in a Desert Rodent Species. Molecular Biology and Evolution, 42(10), msaf230. 10.1093/molbev/msaf230

Yue, L., & Xu, H. (2021). TRP channels in health and disease at a glance. Journal of Cell Science, 134(13), jcs258372. 10.1242/jcs.258372

Yue, Z., Xie, J., Yu, A. S., Stock, J., Du, J., & Yue, L. (2015). Role of TRP channels in the cardiovascular system. American Journal of Physiology. Heart and Circulatory Physiology, 308(3), H157–182. 10.1152/ajpheart.00457.2014

Zakon, H. H. (2012). Adaptive evolution of voltage-gated sodium channels: The first 800 million years. Proceedings of the National Academy of Sciences, 109(supplement_1), 10619-10625. 10.1073/pnas.1201884109

Zakon, H. H., Jost, M. C., & Lu, Y. (2011). Expansion of Voltage-dependent Na+ Channel Gene Family in Early Tetrapods Coincided with the Emergence of Terrestriality and Increased Brain Complexity. Molecular Biology and Evolution, 28(4), 1415–1424. 10.1093/molbev/msq325

Zhang, F., Gu, W., Hurles, M. E., & Lupski, J. R. (2009). Copy Number Variation in Human Health, Disease, and Evolution. Annual Review of Genomics and Human Genetics, 10(1), 451–481. 10.1146/annurev.genom.9.081307.164217

Zhang, G., Li, C., Li, Q., Li, B., Larkin, D. M., Lee, C., Storz, J. F., Antunes, A., Greenwold, M. J., Meredith, R. W., Ödeen, A., Cui, J., Zhou, Q., Xu, L., Pan, H., Wang, Z., Jin, L., Zhang, P., Hu, H., … Froman, D. P. (2014). Comparative genomics reveals insights into avian genome evolution and adaptation. Science, 346(6215), 1311–1320. 10.1126/science.1251385

Zhang, M., Ma, Y., Ye, X., Zhang, N., Pan, L., & Wang, B. (2023). TRP (transient receptor potential) ion channel family: Structures, biological functions and therapeutic interventions for diseases. Signal Transduction and Targeted Therapy, 8(1), 261. 10.1038/s41392-023-01464-x

Zhang, Z., Luo, X., Jiang, L., Wu, H., & Tan, Z. (2024). How do HCN channels play a part in Alzheimer’s and Parkinson’s disease? Ageing Research Reviews, 100, 102436. 10.1016/j.arr.2024.102436

Zhu, F., Feng, M., Sinha, R., Murphy, M. P., Luo, F., Kao, K. S., Szade, K., Seita, J., & Weissman, I. L. (2019). The GABA receptor GABRR1 is expressed on and functional in hematopoietic stem cells and megakaryocyte progenitors. Proceedings of the National Academy of Sciences, 116(37), 18416–18422. 10.1073/pnas.1906251116

